# Cardiac diffusion kurtosis imaging in the human heart in vivo using 300mT/m gradients

**DOI:** 10.1101/2025.01.30.635657

**Authors:** Maryam Afzali, Sam Coveney, Lars Mueller, Sarah Jones, Fabrizio Fasano, C John Evans, Irvin Teh, Erica Dall’Armellina, Filip Szczepankiewicz, Derek K Jones, Jürgen E Schneider

**Author notes:** **Correspondence** Jürgen E Schneider, Biomedical Imaging Science Department, Leeds Institute of Cardiovascular and Metabolic Medicine, University of Leeds, Leeds, LS2 9JT, United Kingdom.

## Abstract

**Purpose:** Diffusion tensor imaging (DTI) is commonly used in cardiac diffusion magnetic resonance imaging (dMRI). However, the tissue’s microstructure (cells, membranes, etc.) restricts the movement of the water molecules, making the spin displacements deviate from Gaussian behaviour. This effect may be observed with diffusion kurtosis imaging (DKI) using sufficiently high b-values (b *>* 450 s*/*mm^2^), which are presently outside the realm of routine cardiac dMRI due to the limited gradient strength of clinical scanners. The Connectom scanner with G_max_ = 300 mT*/*m enables high b-values at echo times (TE) similar to DTI on standard clinical scanners, therefore facilitating cardiac DKI in humans.

**Methods:** Cardiac-gated, second-order motion-compensated dMRI was performed with b_max_ = 1350 s*/*mm^2^ in 10 healthy volunteers on a 3T MRI scanner with G_max_ = 300 mT*/*m. The signal was fitted to a cumulant expansion up to and including the kurtosis term and diffusion metrics such as fractional anisotropy (FA), mean diffusivity (MD), mean kurtosis (MK), axial kurtosis (AK), and radial kurtosis (RK) were calculated.

**Results:** We demonstrate deviation of the signal from monoexponential decay for b-values *>* 450 s*/*mm^2^ (MK = 0.32 *±* 0.03). Radial kurtosis (RK = 0.35 *±* 0.04) was observed slightly larger than axial kurtosis (AK = 0.27 *±* 0.02), and the difference is statistically significant (RK *−* AK = 0.08 *±* 0.04, p = 2e *−* 4).

**Conclusion:** This work demonstrates the feasibility of quantifying kurtosis effect in the human heart in vivo (at an echo time shorter than typical TEs reported for cardiac DTI), using high-performance gradient systems (which are 4-8 times stronger than on standard clinical scanners). Our work lays the foundation for exploring new biomarkers in cardiac dMRI beyond DTI.

## 1 INTRODUCTION

Diffusion magnetic resonance imaging (dMRI) is a non-invasive technique to study tissue microstructure ^1^. So far, diffusion tensor imaging (DTI), which is obtained by combining dMRI measurements along at least six non-collinear encoding directions, has been used most commonly in microstructural investigations of the heart ^2^.

DTI is based on the assumption that the probability of finding a particle in position r at time t adheres to a Gaussian distribution. The standard deviation of this distribution is directly related to the diffusion coefficient. While cardiac diffusion tensor imaging (cDTI) can characterize the average displacement of water molecules in the three-dimensional space, it is not able to provide more specific information about the underlying micro-environment. This is a consequence of the Gaussian assumption in the DTI signal representation ^3^. dMRI signal is usually measured using diffusion-sensitized sequences that vary the diffusion weightings (*b-values*), which are influenced by both the gradient strength and the diffusion time. This signal behavior can be represented using a mono-exponential decay, as described by the Stejskal–Tanner equation ^4^. However, due to cell membranes and other restrictions in biological tissue, the diffusion of water molecules deviates from Gaussian behaviour ^5,6,7^. Consequently, the diffusion-weighted signal in tissue deviates from monoexponential decay at higher b-values as shown for human brain ^8,9^, lung ^10^, prostate ^11^, breast ^12^, calf muscle ^13^, and liver ^14^. However, this has not been widely studied in the human heart in vivo so far.

The DTI representation results from truncating the cumulant expansion of the logarithm of the dMRI signal at the first term where the logarithm of the signal is a linear function of b-value. Conversely, the diffusion kurtosis imaging (DKI) representation truncates at the second term, such that the logarithm of the signal is quadratic in b-value. Diffusional kurtosis is a measure of the restriction of water molecule movement, which in biological tissue is most likely attributable to cell membranes, organelles and tissue compartments, among other factors ^15^. The ability to quantify restriction provides additional microstructural information beyond what is available from the diffusion coefficient alone, making it one of the key advantages of kurtosis imaging ^6^. Tissue structure and other factors, such as the concentration of macromolecules, impact the diffusion coefficient. The diffusion coefficient is therefore a less specific indicator of a tissue’s structural complexity ^6^.

The same type of pulse sequence employed for cardiac diffusion tensor imaging, can be used for DKI, but the required b-values are larger than those usually used to measure diffusion coefficients (b_max_ = 1500 s*/*mm^2^ ^16^). The required b-values can only be obtained at echo times comparable to those used on clinical MR scanners for conventional cDTI ^17,18^ if ultra-strong gradient systems are available / used. The myocardium exhibits significantly shorter T_2_ relaxation times compared to tissues where DTI and DKI are more commonly applied, such as the brain. Consequently, the ability to achieve shorter echo times is paramount for cardiac diffusion imaging. This necessity arises from the need to acquire sufficient signal before T_2_ decay substantially decreases the measurable diffusion-weighted signal, thereby enabling an accurate and reliable diffusion and kurtosis measurements in cardiac tissue.

Non-Gaussian diffusion in the myocardium has been investigated in preclinical experiments such as perfused rat, rabbit, guinea pig and porcine hearts ^19,7,20,21,22,23^. McClymont et al. ^7^ demonstrated non-Gaussian diffusion in healthy and hypertrophic rat hearts and showed that diffusion kurtosis along the second and third eigenvectors of the diffusion tensor can differentiate hypertrophic hearts from sham hearts. Although all of these studies indicate the potential of DKI to provide novel and more refined biomarkers of heart disease, the analysis of diffusion kurtosis in human heart in vivo has been limited. Teh et al. ^16^ have recently demonstrated the feasibility to quantify non-Gaussian diffusion in healthy volunteers using a conventional 3T MR system, albeit at long echo times (TE) (*>* 100 ms) and consequently, low SNR. Furthermore, the reported isotropic, anisotropic and total kurtosis was directionally averaged and not directionally resolved in three dimensions.

Here we investigate three-dimensional diffusion kurtosis in healthy human hearts in vivo using ultra-strong gradients (i.e. 300 mT/m) ^17,18^ at a TE = 61 ms (similar to the echo times commonly used for conventional cardiac DTI) and b_max_ = 1350 s*/*mm^2^ with a second order motion compensated waveform. To the best of our knowledge, this is the first in vivo quantification of three-dimensional diffusion kurtosis in the human heart in vivo.

## 2 THEORY

The apparent kurtosis coefficient for a single direction can be determined by acquiring data at three or more b-values and fitting signal (*S*(*b*)) to the equation:

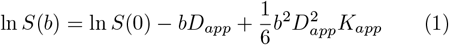

where D_app_ is the apparent diffusion coefficient (ADC) for the given diffusion encoding axis, and K_app_ is the apparent kurtosis coefficient, a dimensionless parameter. The non-Gaussian behavior of water diffusion in the three-dimensional space can be characterized by a symmetric 3 *×* 3 *×* 3 *×* 3 tensor, called the *kurtosis tensor*, W,

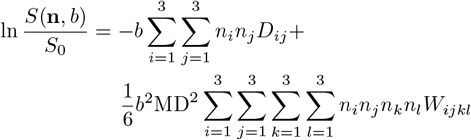

where **n** is the diffusion encoding vector, i, j, k and l are indices of the directions in the physical space, and can take values of 1, 2 or 3. Because of the full symmetry of the tensor, only the following 15 elements ^6^ are independent: W_1111_, *W*_2222_, *W*_3333_, *W*_1112_, *W*_1113_, *W*_1222_, *W*_2223_, *W*_1333_, *W*_2333_, *W*_1122_, *W*_1133_, *W*_2233_, *W*_1123_, *W*_1223_, *W*_1233_.

K_app_ for the direction n = (n_1_, n_2_, n_3_) can be calculated from W_ijkl_:

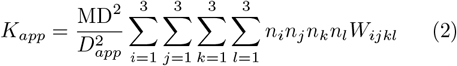

where

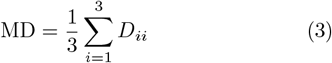

MD is the mean diffusivity and is independent of the direction, and *n*_*i*_ is the component of the direction unit vector.

For DTI, at least 7 measurements with one non-zero b-value is required to quantify the diffusion tensor (6 unique elements due to symmetry) and the non-diffusion weighted signal . We need at least two non-zero b-value and 15 + 6 + 1 = 22 measurements to quantify diffusion tensor D (6 unknown), kurtosis tensor, W (15 unknown), and non-diffusion weighted signal (S_0_). Various diffusion and kurtosis parameters can be calculated from D and W ^24,25,26^, where fractional anisotropy (FA), mean diffusivity (MD), mean kurtosis (MK), axial and radial kurtosis (AK and RK) are amongst the most widely used DKI parameters ^24,27,28^.

## 3 METHODS

### 3.1 Experimental setup and recruitment

Cardiac diffusion-weighted images (cDWI) were acquired on a Connectom 3T research-only MR imaging system (Siemens Healthcare, Erlangen, Germany) with a maximum gradient strength of 300 mT/m and slew rate of 200 T/m/s. An 18-channel body receive coil was used in combination with a 32-channel spine receive coil. Ten healthy volunteers (with no known previous cardiac conditions) were recruited for this study (age range 20.1 *±* 1.6 years (18-22 years), weight range of 64 *±* 12 kg (54-94 kg), six females). The study was approved by the local institutional review board and all subjects provided written consent.

### 3.2 Data acquisition

Routine GRE ^29^ and TRUEFISP ^30^ sequences were used for cardiac planning and cine-imaging, whereas cDWI was performed with a prototype pulse sequence that enabled diffusion encoding with user-defined gradient waveforms ^31,32^. The cine images were acquired in short-axis orientation for apical, mid, and basal slices. cDWI was performed at the same location and orientation as the cine imaging. The phase encoding direction was systematically varied in scout DW images (step size of 30°) and the orientation providing visually the best image quality was chosen for the full cDWI acquisition in each subject.

Diffusion gradient waveforms were designed using the NOW toolbox ^33,34^ (https://github.com/jsjol/NOW) to provide second-order motion compensated waveforms that can reach a specific b-value in the shortest echo time. The maximum gradient strength used in this study for M2-compensation acquisition to generate the b-value of 1350 s*/*mm^2^ was 285.4 mT/m and the maximum slew-rate was 76.2 T/m/s (slew-rate is limited due to peripheral nerve stimulation and cardiac stimulation limits, see ^17^ for more details) which resulted in an echo time of 61 ms (Figure 1). Thus, the waveforms here used the physiologically-limited slew rate of *∼* 76.2 T*/*m*/*s instead of the 200 T/m/s hardware limit. This added 6 ms to the echo time.

**FIGURE 1.**
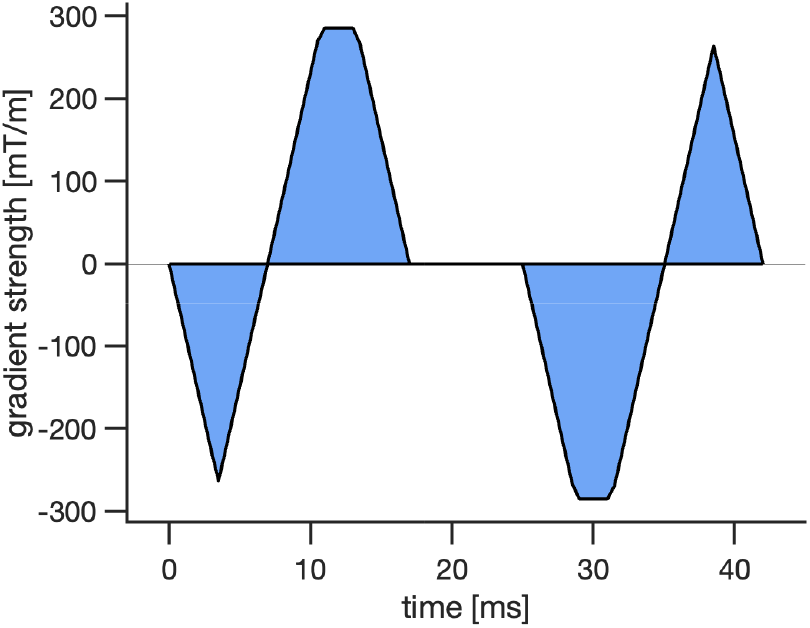
Numerically optimized second-order motion compensated waveform for b_max_ = 1350 s*/*mm^2^, G_max_ = 285.4 mT*/*m and maximum slew rate of 76.2 T*/*m*/*s (lower than the technical limit to reduce PNS and avoid cardiac stimulation).

The cDWI parameters were: TR = 3 RR-intervals, field-of-view = 320 *×* 120 mm^2^ using ZOnally-magnified Oblique Multislice (ZOOM, tilted RF: excitation, tilt angle: 15°, tilted slice thickness: 20 mm) ^32,35^, inplane resolution = 2.7 *×* 2.7 mm^2^, slice thickness = 8 mm, 3 short axis slices (base, mid, and apical), partial Fourier factor = 7/8, no parallel imaging, band-width = 2354 Hz/pixel. Each full data set was comprised of 5 b-values [b = 100, 450, 900, 1200, 1350 s*/*mm^2^] in 30 directions per shell with 6 repetitions, except for the lowest b-value which only had 3 directions and 12 repeats. Data were acquired with ECG-gating and under free-breathing ^2^. Saturation bands were placed around the heart to further suppress the signal from outside the volume of interest. Fat suppression was performed using the SPAIR method ^36^. The trigger delay was adjusted for cDWI acquisition in mid-end systole as determined from the cine images. The total acquisition time was around one hour where the nominal scan time of the DTI/DKI protocol was 40 minutes at 60 beats per minute heart rate. Both magnitude and phase data were collected and used to generate complex-valued images.

### 3.3 Data analysis

The phase variation in each complex-valued diffusion-weighted image was removed using the method proposed by Eichner et al. ^37^. An in-house developed toolbox was used for further post-processing ^38,39^. Real-valued diffusion-weighted images were first registered: for each slice, all low b-value images were registered to one user-specified low b-value image, and then all images were registered to the mean of the co-registered low b-value images. The 2D registration was performed with SimpleITK ^40^, with rigid transformation, separately for basal, middle, and apical slices. Next, an outlier rejection technique was used to semi-automatically remove the outliers (e.g., the images with misregistration or motion corruption) from the data ^41,38^.

The noise level, *σ*, was measured as the standard deviation of the real part of the noise data (acquired without RF pulses) in the image domain from 256 repetitions. The SNR of the data is defined as SNR = S/*σ*, where S is the average of measured signal intensity over the repeats at different b-values in each voxel ^42^. To show the SNR values quantitatively for all subjects, we calculated the SNR per voxel for each b-value and diffusion encoding direction (30 directions for b *>* 100 s*/*mm^2^ and 3 directions for b = 100 s*/*mm^2^), then the SNR values were averaged over different directions for each b-value. The diffusion tensor was fitted to four subsets of data including b = 100 s*/*mm^2^ combined with different maximum b-values (b = 100 and 450 s*/*mm^2^, b = 100 and 900 s*/*mm^2^, b = 100 and 1200 s*/*mm^2^, b = 100 and 1350 s*/*mm^2^) using weighted linear least squares regression (WLS) ^43^, and diffusion metrics such as fractional anisotropy (FA) and mean diffusivity (MD) were calculated for each subset.

The left ventricular myocardium was segmented manually in each slice ^38^. Areas corrupted by susceptibility-related distortion (typically between myocardium, deoxygenated blood, and air, particularly around the posterior vein) were excluded for calculating the global metrics.

Bland–Altman plots were used to compare the mean FA and MD obtained from different subsets of data. To determine the statistical significance between parameters, the Wilcoxon signed-rank test was used; a p-value less than or equal to 0.05 was considered statistically significant.

The DKI representation was then fitted to the data (all b-values) using WLS and the diffusion metrics such as FA, MD, mean kurtosis (MK), axial (AK), and radial kurtosis (RK) were calculated ^44^.

Radial kurtosis (RK) and axial kurtosis (AK) were also compared using Wilcoxon signed-rank test.

## 4 RESULTS

Figure 2 shows representative diffusion-weighted images averaged over six repeats of a single diffusion direction acquired with b = 100, 450, 900, 1200, and 1350 s*/*mm^2^ in short axis view. On average 12% *±* 6% of diffusion-weighted images were discarded due to poor image quality and signal dropout.

**FIGURE 2.**
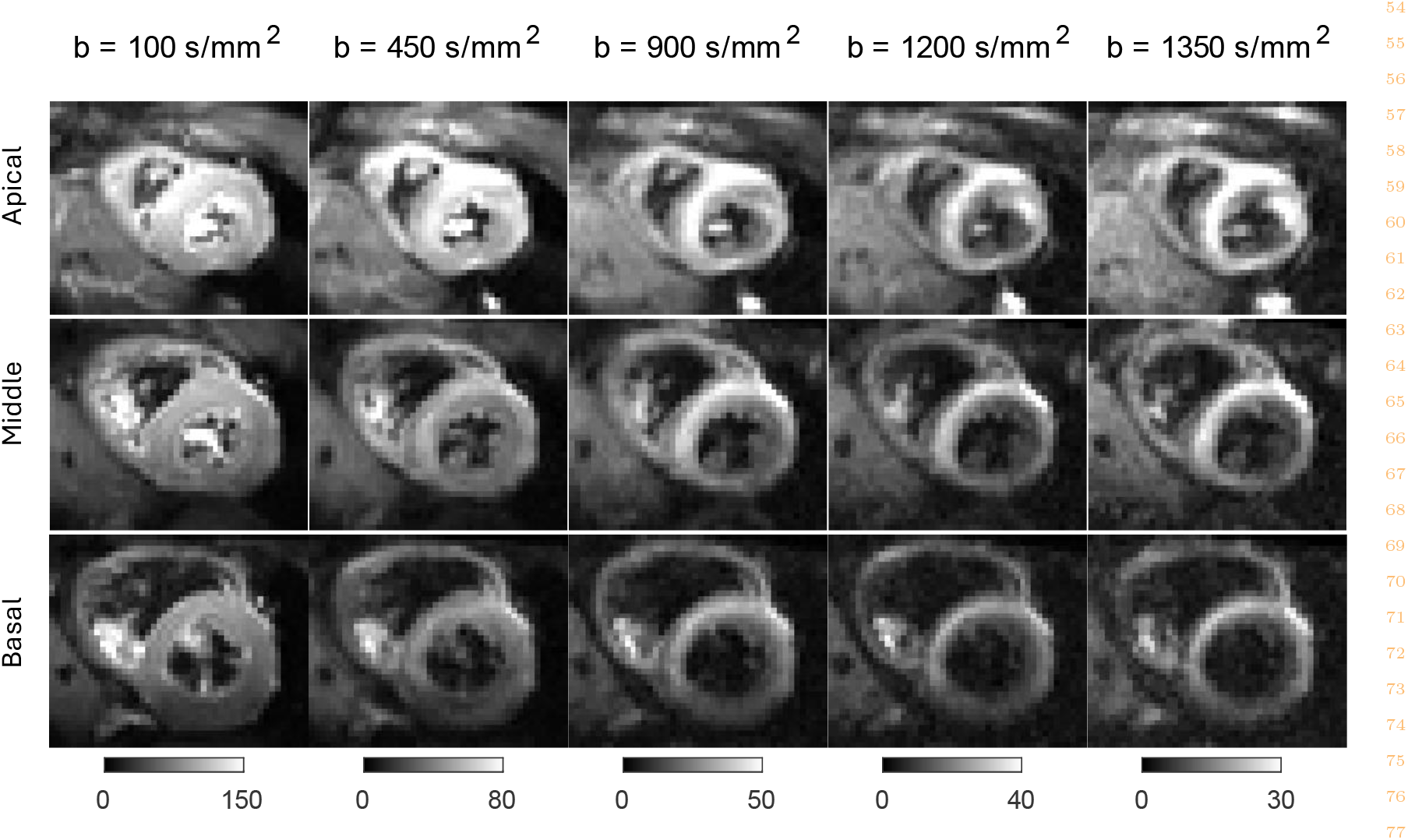
Example representative cardiac diffusion-weighted images averaged over six repeats of a single diffusion direction acquired in basal, middle, and apical slices with b-value = 100, 450, 900, 1200 and 1350 s*/*mm^2^ using second-order motion compensation with TE = 61 ms.

The measured SNR of the diffusion weighted images over the left ventricle for an arbitrary orientation at b = 100, 450, 900, 1200, 1350 s*/*mm^2^ is shown in Figure 3 for a single subject. Each b-value uses a different window/level for better visibility. The mean and standard deviation of the SNR at b = 100, 450, 900, 1200, and 1350 s*/*mm^2^ were 40 *±* 10, 23 *±* 6, 12 *±* 3, 9 *±* 2, 7 *±* 2, respectively.

**FIGURE 3.**
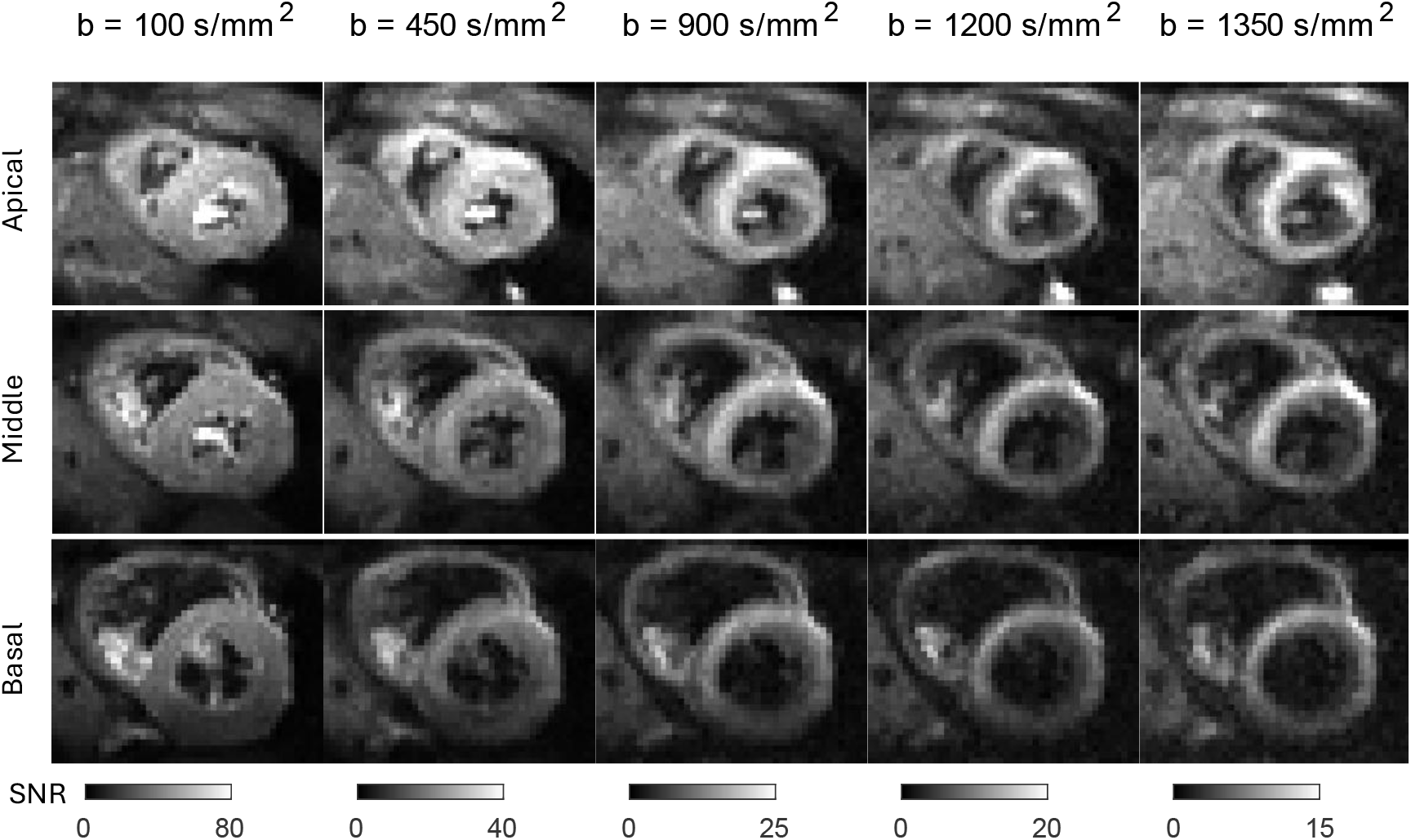
Signal-to-noise-ratio (SNR) maps obtained in basal, middle, and apical slices from a single diffusion direction (the same diffusion direction as Figure 2) with b = 100, 450, 900, 1200 and 1350 s*/*mm^2^ using second order motion compensation (M_2_, TE = 61 ms).

The mean MD and FA values obtained from DTI fitting of four subsets of data with different maximum b-values (b = 100 and 450 s*/*mm^2^, b = 100 and 900 s*/*mm^2^, b = 100 and 1200 s*/*mm^2^, b = 100 and 1350 s*/*mm^2^) for 10 subjects are shown as boxplots in Figure 4 . The average MD value reduces from 1.58 *±* 0.04 *×* 10^*−*3^mm^2^*/*s at b_max_ = 450 s*/*mm^2^ to 1.44 *±* 0.04 *×* 10^*−*3^mm^2^*/*s at b_max_ = 1350 s*/*mm^2^, and the difference is statistically significant (p *<* 0.05) (Table 2). While the mean FA values are almost unchanged at different b_max_ values (0.31 *±* 0.01).

**TABLE 1.** Mean *±* standard deviation of mean diffusivity (MD) and fractional anisotropy (FA) for DTI fit to different b_max_ (b_max_ = 450 s*/*mm^2^ [b = 100 and 450 s*/*mm^2^], b_max_ = 900 s*/*mm^2^ [b = 100 and 900 s*/*mm^2^], b_max_ = 1200 s*/*mm^2^ [b = 100 and 1200 s*/*mm^2^], b_max_ = 1350 s*/*mm^2^ [b = 100 and 1350 s*/*mm^2^]) inside a left ventricle mask and then averaged over volunteers.

| $b_{\max}$ [s/mm<sup>2</sup>] | MD [$\times 10^{-3}$ mm<sup>2</sup>/s] | FA |
| --- | --- | --- |
| 450 | 1.58 ± 0.04 | 0.30 ± 0.01 |
| 900 | 1.51 ± 0.04 | 0.31 ± 0.01 |
| 1200 | 1.46 ± 0.04 | 0.31 ± 0.01 |
| 1350 | 1.44 ± 0.04 | 0.31 ± 0.01 |

**TABLE 2.** Mean difference *±* standard deviation between global mean values of MD, FA calculated using DTI fit to b_450_ : b = 100, and 450 s*/*mm^2^ compared to b_900_ : b = 100, and 900 s*/*mm^2^, b_1200_ : b = 100, and 1200 s*/*mm^2^, and b_1350_ : b = 100, and 1350 s*/*mm^2^ inside a left ventricle mask.

|  |  |
| --- | --- |
| MD($b_{900}$) - MD($b_{450}$) [$\times 10^{-3} \text{ mm}^2/\text{s}$] | $0.07 \pm 0.02$ ($p = 2\text{e-}7$) (Figure 4 (B)) |
| MD($b_{1200}$) - MD($b_{450}$) [$\times 10^{-3} \text{ mm}^2/\text{s}$] | $0.12 \pm 0.01$ ($p = 5\text{e-}10$) (Figure 4 (C)) |
| MD($b_{1350}$) - MD($b_{450}$) [$\times 10^{-3} \text{ mm}^2/\text{s}$] | $0.14 \pm 0.01$ ($p = 2\text{e-}10$) (Figure 4 (D)) |
| FA($b_{900}$) - FA($b_{450}$) | $-0.001 \pm 0.005$ ($p = 0.37$) (Figure 4 (F)) |
| FA($b_{1200}$) - FA($b_{450}$) | $-0.005 \pm 0.007$ ($p = 0.03$) (Figure 4 (G)) |
| FA($b_{1350}$) - FA($b_{450}$) | $-0.005 \pm 0.008$ ($p = 0.06$) (Figure 4 (H)) |

**FIGURE 4.**
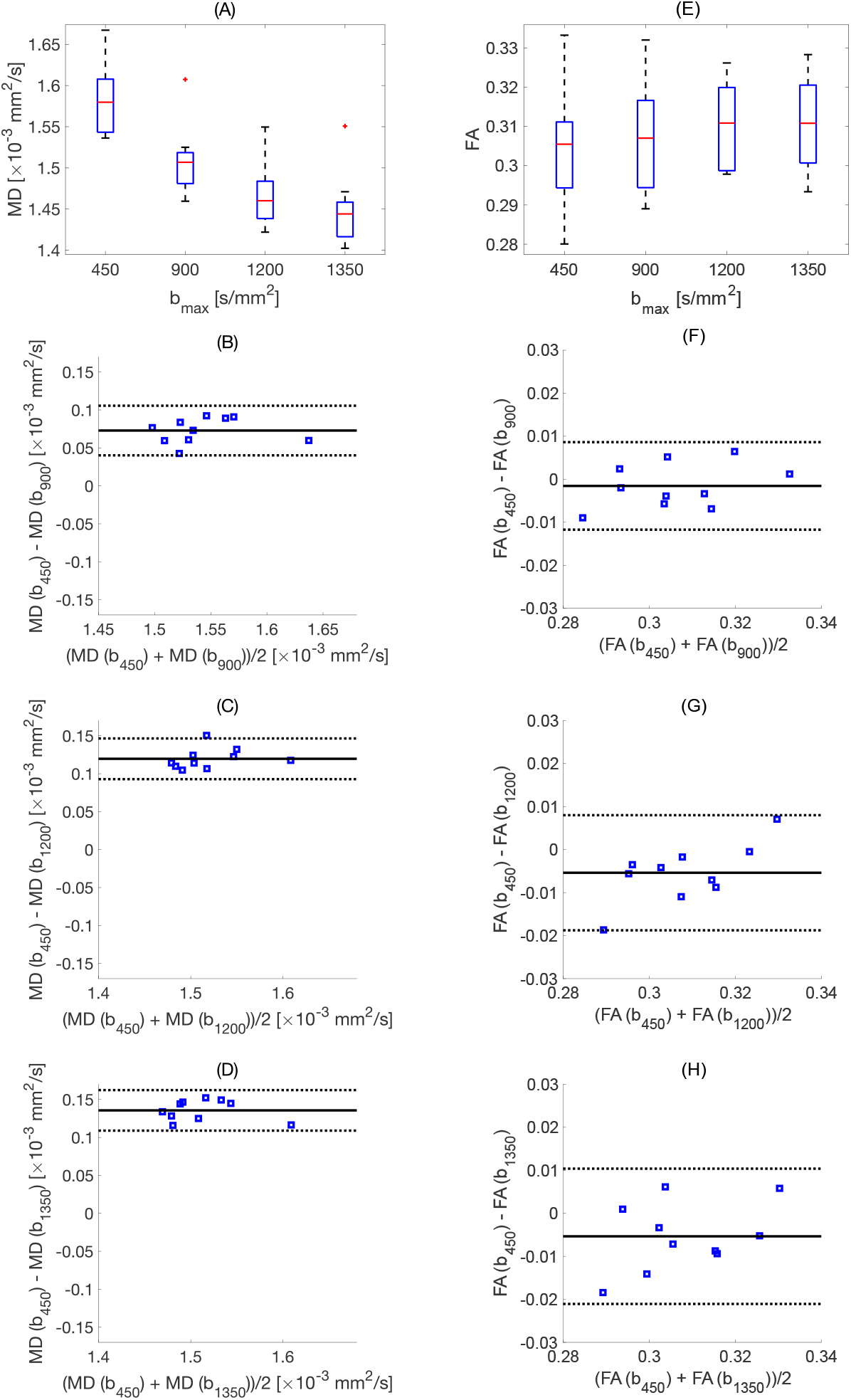
Change of MD and FA for different b_max_ values. (A) and (E) show the box plot of mean MD and FA over 10 subjects (the red line shows the mean value). (B-D) and (F-H) Bland–Altman plots, comparing mean diffusivity (MD), and fractional anisotropy (FA), calculated using DTI fit to b_450_ : b = 100, and 450 s*/*mm^2^ compared to (B) and (F) b_900_ : b = 100, and 900 s*/*mm^2^, (C) and (G) b_1200_ : b = 100, and 1200 s*/*mm^2^, and (D) and (H) b_1350_ : b = 100, and 1350 s*/*mm^2^. Mean difference *±* 1.96 SD is given by solid and dashed black lines, respectively (N = 10 subjects).

Figure 5 shows the average signal attenuation versus b-value in the ROI highlighted as green for a randomly chosen subject. The blue dots and error bars indicate the mean and standard deviation of the measured signal over the ROI at each b-value while the mono-exponential fit to the averaged signal from b *≤* 450 s*/*mm^2^ is shown in red and the DKI fit in yellow. Note that, here we use a one-dimensional mono-exponential fit (red curve) and one-dimensional DKI as described in Eq. 1 (yellow curve). At lower b-values, b *≤* 450 s*/*mm^2^, the signal from the mono-exponential fit and the measured signal are indistinguishable, as expected from the theory. However, the measured signal clearly deviates from the mono-exponential decay for b *>* 450 s*/*mm^2^. The relative difference between mono-exponential fit and the measured signal generally increases with b-value, corresponding to the divergent signal attenuation curves, which is a hallmark of non-Gaussian diffusion in the tissue.

**FIGURE 5.**
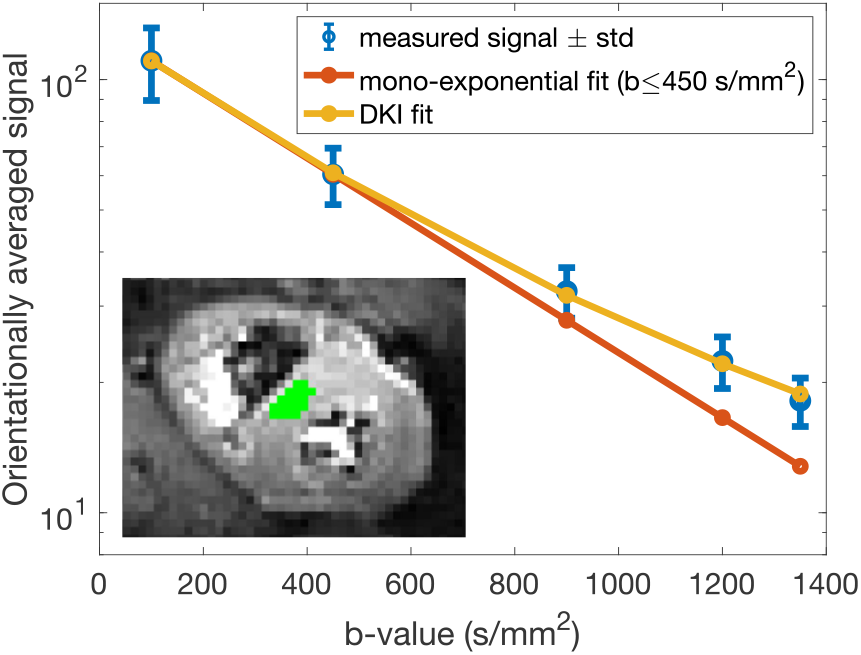
Semi-log plot of the average signal over the region of interest (ROI), highlighted as green, for each b-value. The blue dots and the error bars show the mean and standard deviation of the measured signal in the ROI, the red curve shows the mono-exponential fit to the average signal from b *≤* 450 s*/*mm^2^, and the yellow curve shows the one-dimensional DKI fit. The measured signal deviates from the mono-exponential decay by increasing the b-value above 450 s*/*mm^2^.

Representative FA, MD, HA, and E2A maps from DTI and FA, MD, HA, E2A, MK, AK, and RK from DKI fitting for apical, middle and basal slices are shown in Figure 6 .

**FIGURE 6.**
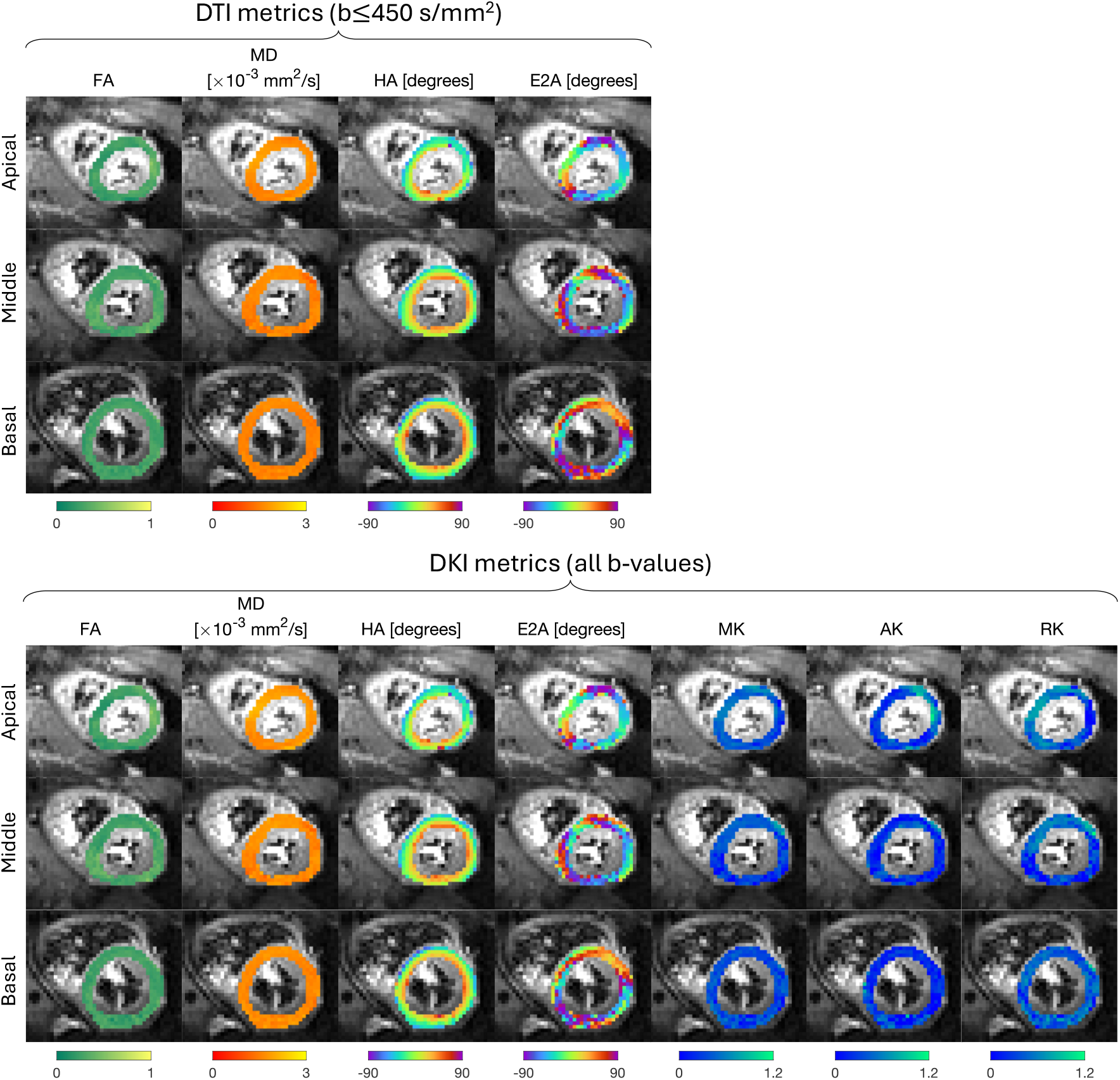
Examples of mean diffusivity (MD), fractional anisotropy (FA), helix angle (HA) and secondary eigenvector angle (E2A) calculated using b = 100 and 450 s*/*mm^2^ for DTI and FA, MD, HA, E2A, mean, axial and radial kurtosis (MK, AK, and RK) calculated using all b-values for DKI.

The mean and standard deviation of MD, FA, MK, AK, and RK for all 10 subjects are shown in Figure 7 . The average value of the DTI and DKI metrics over 10 subjects are presented in Table 3 (DTI: MD = 1.58 *±* 0.04 s*/*mm^2^ and FA = 0.30 *±* 0.01. DKI: MD = 1.66 *±* 0.04, FA = 0.31 *±* 0.02, MK = 0.32 *±* 0.03, AK = 0.27 *±* 0.02, and RK = 0.35 *±* 0.04). It was found that the radial kurtosis (RK) was slightly higher than axial kurtosis (AK) (RK = 0.35 *±* 0.06 vs. AK = 0.27 *±* 0.05) and the difference is statistically significant (RK *−* AK = 0.08 *±* 0.04 (p = 2 *×* 10^*−*4^)) (Table 3 and Figure 7 (D)).

**TABLE 3.** Mean *±* standard deviation of mean diffusivity (MD) and fractional anisotropy (FA) for DTI (b = 100 and 450 s*/*mm^2^) and MD, FA, mean kurtosis (MK), axial kurtosis (AK), and radial kurtosis (RK) for DKI (all b-values) inside a left ventricle mask and then averaged over volunteers.

| DTI | MD [$\times 10^{-3} \text{ mm}^2/\text{s}$] | FA | | | |
| --- | --- | --- | --- | --- | --- |
| | $1.58 \pm 0.04$ | $0.30 \pm 0.01$ | | | |
| DKI | MD [$\times 10^{-3} \text{ mm}^2/\text{s}$] | FA | MK | AK | RK |
| | $1.66 \pm 0.04$ | $0.31 \pm 0.02$ | $0.32 \pm 0.03$ | $0.27 \pm 0.02$ | $0.35 \pm 0.04$ |

**FIGURE 7.**
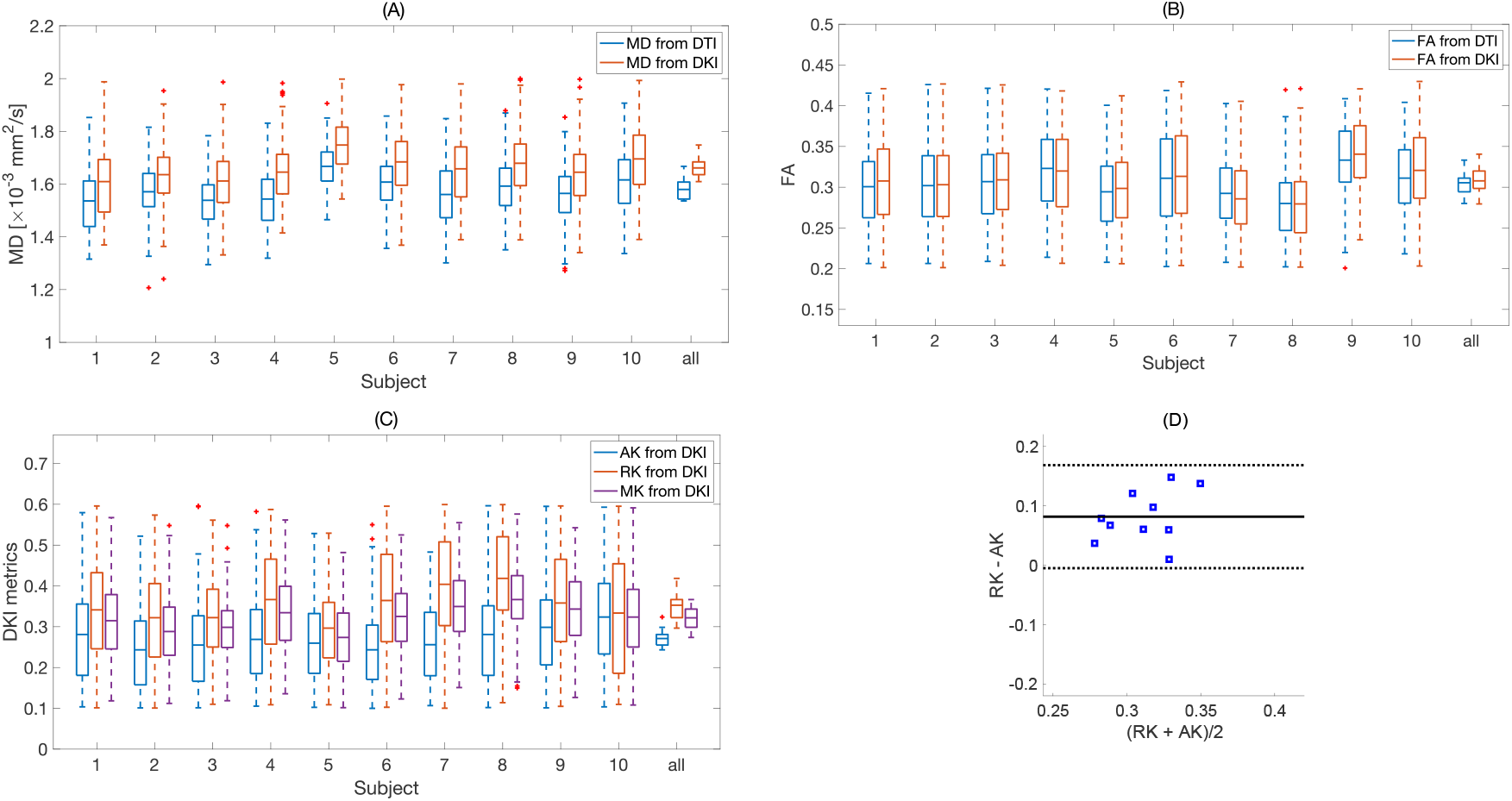
Box plot of (A) mean diffusivity (MD), (B) fractional anisotropy (FA) from DTI and DKI, (C) mean kurtosis (MK), axial kurtosis (AK), and radial kurtosis from DKI over left ventricular mask for all 10 subjects (“all” represents the distribution of the mean values in subjects 1 to 10). On each box, the central mark indicates the mean, and the bottom and top edges of the box indicate the 25th and 75th percentiles, respectively. The whiskers extend to the most extreme data points not considered outliers, and the outliers are plotted individually using the ‘+’ marker symbol. (D) Bland–Altman plot, comparing axial kurtosis (AK) and radial kurtosis (RK) from DKI. Mean difference *±* 1.96 SD is given by solid and dashed black lines, respectively (N = 10 subjects).

## 5 DISCUSSION

This work demonstrates the feasibility of performing DKI in the human heart in vivo using much stronger gradients than commonly available in routine clinical settings. To accurately measure diffusional kurtosis, b-values higher than those typically used in cardiac DTI are necessary to clearly reveal the deviation from mono-exponential decay ^7,18,16^.

In brain studies, maximum b-values of about 2000 to 3000 s*/*mm^2^ ^24^ (two to three times higher than b = 1000 s*/*mm^2^ used for DTI) are recommended to quantify the kurtosis effect. We followed the same rational, the maximum b-value used in cardiac DTI in vivo is around 450 s*/*mm^2^, so the maximum b-value of 900 to 1350 s*/*mm^2^ should be sufficiently high to show the deviation of diffusion weighted signal from the mono-exponential decay. Sufficient SNR at high b-value images is essential to accurately estimate DKI metrics ^45^. Glenn et al. ^46^ found that the estimated kurtosis parameters are 93% accurate when the SNR is greater than 3 and greater than 98% for SNR greater than 5. For insufficient SNR, the signal intensity approaches the noise floor, resulting in an artificial curvature of the signal decay and biased estimates of the kurtosis metrics ^46^. Tissue diffusion properties define the rate at which signal approaches the noise floor. Obtaining sufficient SNR at high b-values is particularly challenging in the human heart due to motion and the associated need for motion-compensated gradient waveforms, which in turn require longer echo times, additionally confounded by short T_2_ of the heart tissue ^47^. The use of high-performance gradient system in this study, a b-value of 1350 s*/*mm^2^ with a minimum TE = 61 ms for the chosen imaging parameters and optimized waveforms was feasible, which significantly improved the SNR efficiency of in vivo cardiac diffusion kurtosis imaging imaging. The achieved sequence timings were comparable to standard cardiac DTI acquisitions at more than three times (*>* 3*×*) higher b-values. It provides SNR of 7 *±* 2 over the left ventricle for b = 1350 s*/*mm^2^(Figure 3) which is high enough to avoid diffusion weighted signal falling below the noise floor ^46^. Notably, the same b-value on clinical routine systems with G_max_= 80 mT/m would need an echo time of at least 90 ms (Teh et al. ^16^ used TE = 118 ms to enable tensor-valued diffusion encoding at b-value of 1500 s*/*mm^2^). The approximately 30 ms shorter echo time improves the SNR nearly two-fold due to the short T_2_ of cardiac tissue: based on the reported T_2_ of around 46 ms ^48^ the SNR increase is exp(*−*61*/*46)*/* exp(*−*90*/*46) *≈* 1.88.

We found that by increasing b-value, the MD values from DTI fit decreased (MD = 1.58 *±* 0.04 *×* 10^*−*3^mm^2^*/*s at b_m_ax450 s*/*mm^2^ and MD = 1.44 *±* 0.04 *×* 10^*−*3^mm^2^*/*s at b_m_ax1350 s*/*mm^2^) where this difference is statistically significant (p = 2e *−* 10) (Figure 4, Table 1, 2). This reduction in MD is due to the kurtosis effect that is more pronounced at higher b-values. One may argue that the noise in the data might have a similar effect, however the SNR at the highest b-value is 7 *±* 2 and also the real part of the signal (instead of magnitude) is used in this work to avoid the bias in the signal due to rician noise ^37^, therefore the reduction in MD should be due to the kurtosis effect.

To the best of our knowledge, this is the first report of three-dimensional diffusion kurtosis in the human heart in vivo. We found that radial kurtosis (RK = 0.35 *±* 0.04) was slightly larger than axial kurtosis (AK = 0.27 *±* 0.02) and the difference was statistically significant (RK - AK = 0.08 *±* 0.04, p = 2e *−* 4, Table 3 and Figure 7 (D)). Teh et al. ^16^ previously reported total kurtosis of 0.33 *±* 0.09 for in vivo human heart on a standard clinical MR scanner which aligns with our results on ultra-strong gradient system (MK = 0.32 *±* 0.03), however they did not quantify the three-dimensional kurtosis metrics such as AK and RK. It is known that there are more restrictions perpendicular to the cardiomyocyte orientations than parallel orientations and previous study on ex vivo rat ^7^ and pig hearts ^23^ showed higher RK compared to AK. Mcclymont et al. ^7^ reported a kurtosis value of 0.13 *±* 0.02, 0.45 *±* 0.04 and 0.55 *±* 0.03 along first, second and third eigenvector. We observe the trend of higher RK compared to AK in our work, however the kurtosis values in the in vivo human heart are slightly different than the values reported ex vivo. The reason is that the tissue preparation / fixation may affect the cell size and cause shrinkage which results in higher kurtosis effect along the perpendicular direction. In addition the difference between species, i.e. rat heart versus human heart may contribute to this difference as well.

The magnitude of kurtosis observed in the in vivo human heart is limited due to the microstructure of the cardiac tissue. We used a simple biophyscial model to simulate the signal from cardiac tissue and the estimated kurtosis values in the simulated signal are close to the values we obtained in vivo (see Supplementary information for more details). We further investigated the effect of diffusion encoding time on the kurtosis values and we found that by increasing the effective diffusion time, the amount of mean and radial kurtosis (MK and RK) slightly increase (Table S1), however, the increase in diffusion encoding time, prolonges the echo time and therefore reduces the SNR.

The study was conducted on a limited cohort of healthy volunteers, extrapolation of this technique to patient populations with specific cardiac pathologies is the topic of our future work. Diffusion kurtosis metrics are sensitive to tissue microstructure but are not specific to particular microstructural features (e.g., intracelluar and extracellular space).

The primary purpose of our work was to establish the feasibility of DKI in human hearts in vivo using ultra-strong gradients and highly relevant with the advent of the latest generation of clinical MR scanners with 200 mT/m gradient systems (such as the Siemens Healthineers, Magnetom Cima.X). This opens the field for novel investigation and paves the way for the exploration of cardiac DKI in clinical studies.

## 6 CONCLUSION

In this work, we demonstrated the feasibility of three-dimensional DKI in the human heart in vivo. This was facilitated using strong gradients (300 mT/m) that could provide high b-values (b_max_ = 1350 s*/*mm^2^) for a second order motion compensated waveform at a TE = 61 ms. Radial kurtosis (RK) was observed slightly larger than axial kurtosis (AK) as expected and the difference was statistically significant. The in vivo measurement of radial, axial and mean kurtosis provides the potential for characterizing the myocardial microstructure, which may be useful in some cardiac diseases.

## 7 ACKNOWLEDGMENTS

This research was funded in whole, or in part, by Wellcome Trust Investigator Award (219536/Z/19/Z) and (096646/Z/11/Z), a Wellcome Trust Strategic Award (104943/Z/14/Z), the EPSRC (EP/M029778/1), The Wolfson Foundation, and the British Heart Foundation (PG/19/1/34076) and Swedish Cancer Society (22 0592 JIA). For the purpose of open access, the author has applied a CC BY public copyright license to any Author Accepted Manuscript version arising from this submission. We thank Siemens Healthineers for the pulse sequence development environment.

## Author contributions

Study conceptualization, design, and planning (Maryam Afzali, Jürgen E Schneider, Lars Mueller, Sam Coveney, Derek K Jones, Irvin Teh), funding (Jürgen E Schneider), MRI pulse sequence design (Filip Szczepankiewicz, Lars Mueller, Fabrizio Fasano), experimental design (Maryam Afzali, Sam Coveney, Jürgen E Schneider, Lars Mueller), MRI data acquisition (Maryam Afzali, Lars Mueller, Sarah Jones, Erica Dall’Armellina, Fabrizio Fasano, John Evans), MRI data analysis (Maryam Afzali, Sam Coveney), clinical expertise (Erica Dall’Armellina), ethics approvals (Maryam Afzali, Jürgen E Schneider, John Evans), drafting of manuscript (Maryam Afzali), and manuscript review (All).

## REFERENCES

1. Moseley ME, Kucharczyk J, Mintorovitch J, et al. Diffusion-weighted MR imaging of acute stroke: correlation with T2-weighted and magnetic susceptibility-enhanced MR imaging in cats.. American Journal of Neuroradiology. 1990;11(3):423–429.

2. Das Arka, Kelly Christopher, Teh Irvin, et al. Phenotyping hypertrophic cardiomyopathy using cardiac diffusion magnetic resonance imaging: the relationship between microvascular dysfunction and microstructural changes. European Heart Journal-Cardiovascular Imaging. 2022;23(3):352–362.

3. Basser Peter J, Mattiello James, LeBihan Denis. MR diffusion tensor spectroscopy and imaging. Biophysical journal. 1994;66(1):259–267.

4. Stejskal Edward O, Tanner John E. Spin diffusion measurements: spin echoes in the presence of a time-dependent field gradient. The journal of chemical physics. 1965;42(1):288–292.

5. Alexander DC, Barker GJ, Arridge SR. Detection and modeling of non-Gaussian apparent diffusion coefficient profiles in human brain data. Magnetic Resonance in Medicine: An Official Journal of the International Society for Magnetic Resonance in Medicine. 2002;48(2):331– 340.

6. Jensen Jens H, Helpern Joseph A, Ramani Anita, Lu Hanzhang, Kaczynski Kyle. Diffusional kurtosis imaging: the quantification of non-Gaussian water diffusion by means of magnetic resonance imaging. Magnetic Resonance in Medicine: An Official Journal of the International Society for Magnetic Resonance in Medicine. 2005;53(6):1432–1440.

7. McClymont Darryl, Teh Irvin, Carruth Eric, et al. Evaluation of non-Gaussian diffusion in cardiac MRI. Magnetic resonance in medicine. 2017;78(3):1174–1186.

8. Falangola Maria F, Jensen Jens H, Babb James S, et al. Age-related non-Gaussian diffusion patterns in the prefrontal brain. Journal of Magnetic Resonance Imaging: An Official Journal of the International Society for Magnetic Resonance in Medicine. 2008;28(6):1345–1350.

9. Afzali Maryam, Pieciak Tomasz, Jones Derek K, Schneider Jürgen E, Özarslan Evren. Cumulant Expansion with Localization: A new representation of the diffusion MRI signal. Frontiers in Neuroimaging. 2022;1:958680.

10. Trampel Robert, Jensen Jens H, Lee Ray F, Kamenetskiy Igor, McGuinness Georgeann, Johnson Glyn. Diffusional kurtosis imaging in the lung using hyperpolarized 3He. Magnetic Resonance in Medicine: An Official Journal of the International Society for Magnetic Resonance in Medicine. 2006;56(4):733–737.

11. Rosenkrantz Andrew B, Sigmund Eric E, Johnson Glyn, et al. Prostate cancer: feasibility and preliminary experience of a diffusional kurtosis model for detection and assessment of aggressiveness of peripheral zone cancer. Radiology. 2012;264(1):126–135.

12. Wu Dongmei, Li Guanwu, Zhang Junxiang, Chang Shixing, Hu Jiani, Dai Yongming. Characterization of breast tumors using diffusion kurtosis imaging (DKI). PloS one. 2014;9(11):e113240.

13. Marschar Anja Maria, Kuder Tristan Anselm, Stieltjes Bram, Nagel Armin Michael, Bachert Peter, Laun Frederik Bernd. In vivo imaging of the time-dependent apparent diffusional kurtosis in the human calf muscle. Journal of Magnetic Resonance Imaging. 2015;41(6):1581–1590.

14. Rosenkrantz Andrew B, Sigmund Eric E, Winnick Aaron, et al. Assessment of hepatocellular carcinoma using apparent diffusion coefficient and diffusion kurtosis indices: preliminary experience in fresh liver explants. Magnetic resonance imaging. 2012;30(10):1534–1540.

15. Hui Edward S, Du Fang, Huang Shiliang, Shen Qiang, Duong Timothy Q. Spatiotemporal dynamics of diffusional kurtosis, mean diffusivity and perfusion changes in experimental stroke. Brain research. 2012;1451:100–109.

16. Teh Irvin, Shelley David, Boyle Jordan H, et al. Cardiac q-space trajectory imaging by motion-compensated tensor-valued diffusion encoding in human heart in vivo. Magnetic Resonance in Medicine. 2023;90(1):150–165.

17. Afzali Maryam, Mueller Lars, Coveney Sam, et al. In vivo diffusion MRI of the human heart using a 300 mT/m gradient system. Magnetic Resonance in Medicine. 2024;.

18. Afzali Maryam, Mueller L, Coveney C, et al. Quantification of non-Gaussian diffusion in the human heart in vivo. In: ; 2024.

19. Hsu Edward W, Buckley David L, Bui Jonathan D, Blackband Stephen J, Forder John R. Two-component diffusion tensor MRI of isolated perfused hearts. Magnetic Resonance in Medicine: An Official Journal of the International Society for Magnetic Resonance in Medicine. 2001;45(6):1039–1045.

20. Wu Yin, Zou Chao, Liu Wei, et al. Effect of B-value in revealing postinfarct myocardial microstructural remodeling using MR diffusion tensor imaging. Magnetic resonance imaging. 2013;31(6):847–856.

21. Forder John R, Bui Jonathan D, Buckley David L, Blackband Stephen J. MR imaging measurement of compartmental water diffusion in perfused heart slices. American Journal of Physiology-Heart and Circulatory Physiology. 2001;281(3):H1280–H1285.

22. Abdullah Osama M, Gomez Arnold David, Merchant Samer, Heidinger Michael, Poelzing Steven, Hsu Edward W. Orientation dependence of microcirculation-induced diffusion signal in anisotropic tissues. Magnetic resonance in medicine. 2016;76(4):1252–1262.

23. Mazzoli Valentina, Froeling Martijn, Nederveen Aart J, Nicolay Klaas, Strijkers Gustav J. Cardiac diffusion MRI beyond DTI. In: ; 2014.

24. Jensen Jens H., Helpern Joseph A.. MRI quantification of non-Gaussian water diffusion by kurtosis analysis. NMR in Biomedicine. 2010;23(7):698–710.

25. Tabesh Ali, Jensen Jens H, Ardekani Babak A, Helpern Joseph A. Estimation of tensors and tensor-derived measures in diffusional kurtosis imaging. Magnetic resonance in medicine. 2011;65(3):823–836.

26. Glenn G Russell, Helpern Joseph A, Tabesh Ali, Jensen Jens H. Quantitative assessment of diffusional kurtosis anisotropy. NMR in Biomedicine. 2015;28(4):448–459.

27. Henriques Rafael Neto, Correia Marta Morgado, Nunes Rita Gouveia, Ferreira Hugo Alexandre. Exploring the 3D geometry of the diffusion kurtosis tensor—Impact on the development of robust tractography procedures and novel biomarkers. Neuroimage. 2015;111:85–99.

28. Henriques Rafael N, Jespersen Sune N, Jones Derek K, Veraart Jelle. Toward more robust and reproducible diffusion kurtosis imaging. Magnetic resonance in medicine. 2021;86(3):1600–1613.

29. Winkler Mark L, Ortendahl DA, Mills TC, et al. Characteristics of partial flip angle and gradient reversal MR imaging.. Radiology. 1988;166(1):17–26.

30. Oppelt A. FISP: eine neue schnelle Pulssequenz für die Kernspintomographie. Electromedica. 1986;54:15–18.

31. Szczepankiewicz Filip, Sjölund Jens, Ståhlberg Freddy, Lätt Jimmy, Nilsson Markus. Tensor-valued diffusion encoding for diffusional variance decomposition (DIVIDE): Technical feasibility in clinical MRI systems. PLoS One. 2019;14(3):e0214238.

32. Mueller Lars, Afzali M, Coveney S, et al. ZOOM and enhance: ZOnally magnified Oblique Multi-slice for cardiac DTI with ultra-strong gradients. In: ; 2024.

33. SjÖlund Jens, Szczepankiewicz Filip, Nilsson Markus, Topgaard Daniel, Westin Carl-Fredrik, Knutsson Hans. Constrained optimization of gradient waveforms for generalized diffusion encoding. Journal of Magnetic Resonance. 2015;261:157–168.

34. Szczepankiewicz Filip, Sjölund Jens, Dall’Armellina Erica, et al. Motion-compensated gradient waveforms for tensor-valued diffusion encoding by constrained numerical optimization. Magnetic resonance in medicine. 2021;85(4):2117–2126.

35. Symms MR, Wheeler-Kingshott CA, Parker GJM, Barker GJ. Zonally-magnified oblique multislice (ZOOM) EPI. In: ; 2000.

36. Lauenstein Thomas C, Sharma Puneet, Hughes Timothy, Heberlein Keith, Tudorascu Dana, Martin Diego R. Evaluation of optimized inversion-recovery fat-suppression techniques for T2-weighted abdominal MR imaging. Journal of Magnetic Resonance Imaging. 2008;27(6):1448–1454.

37. Eichner Cornelius, Cauley Stephen F, Cohen-Adad Julien, et al. Real diffusion-weighted MRI enabling true signal averaging and increased diffusion contrast. NeuroImage. 2015;122:373–384.

38. Coveney C, Kelly C, Teh I, et al. Semi-automated rejection of corrupted images in cardiac diffusion tensor imaging. Proceedings of the Annual Meeting of ISMRM. 2023;.

39. Garyfallidis Eleftherios, Brett Matthew, Amirbekian Bagrat, et al. Dipy, a library for the analysis of diffusion MRI data. Frontiers in neuroinformatics. 2014;8:8.

40. Lowekamp Bradley C, Chen David T, Ibáñez Luis, Blezek Daniel. The design of SimpleITK. Frontiers in neuroinformatics. 2013;7:45.

41. Coveney Sam, Afzali Maryam, Mueller Lars, et al. Outlier detection in cardiac diffusion tensor imaging: Shot rejection or robust fitting?. Medical Image Analysis. 2025;101:103386.

42. McClymont Darryl, Teh Irvin, Schneider Jürgen E. The impact of signal-to-noise ratio, diffusion-weighted directions and image resolution in cardiac diffusion tensor imaging–insights from the ex-vivo rat heart. Journal of Cardiovascular Magnetic Resonance. 2017;19(1):1–10.

43. Salvador Raymond, Peña Alonso, Menon David K, Carpenter T Adrian, Pickard John D, Bullmore Ed T. Formal characterization and extension of the linearized diffusion tensor model. Human brain mapping. 2005;24(2):144– 155.

44. Hansen Brian, Shemesh Noam, Jespersen Sune Nørhøj. Fast imaging of mean, axial and radial diffusion kurtosis. Neuroimage. 2016;142:381–393.

45. Lu Hanzhang, Jensen Jens H, Ramani Anita, Helpern Joseph A. Three-dimensional characterization of nongaussian water diffusion in humans using diffusion kurtosis imaging. NMR in Biomedicine: An International Journal Devoted to the Development and Application of Magnetic Resonance In vivo. 2006;19(2):236–247.

46. Glenn G Russell, Tabesh Ali, Jensen Jens H. A simple noise correction scheme for diffusional kurtosis imaging. Magnetic resonance imaging. 2015;33(1):124–133.

47. Scott Andrew D, Ferreira Pedro Fadc, Nielles-Vallespin Sonia, et al. Optimal diffusion weighting for in vivo cardiac diffusion tensor imaging. Magnetic resonance in medicine. 2015;74(2):420–430.

48. Hanson Christopher A, Kamath Akshay, Gottbrecht Matthew, Ibrahim Sami, Salerno Michael. T2 relaxation times at cardiac MRI in healthy adults: a systematic review and meta-analysis. Radiology. 2020;297(2):344– 351.

