## supporting information for "Cardiac diffusion kurtosis imaging in the human heart in vivo using 300mT/m gradients"

Cardiomyocytes are 17 to 25 $\mu$m in diameter (Tracy and Sander, 2011) therefore the amount of restriction is limited compared to the brain white matter where the axons have a radius of a few micrometers. To support this statement, we simulated the signal from a simple multi-compartment model including a cylinder representative of cardiomyocyte and a zeppelin for the extra-cellular space. Therefore, the simulated diffusion-weighted signal is described as a sum of two components:

$$\mathrm{S} = \mathrm{f_{cylinder}} \mathrm{S_{cylinder}}(\mathrm{R_{cylinder}}, \mathrm{D^{||}_{cylinder}}) + \mathrm{f_{zeppelin}} \mathrm{S_{zeppelin}}(\mathrm{D^{||}_{zeppelin}}, \mathrm{D^{\perp}_{zeppelin}}) \tag{R1}$$

where $\mathrm{f_{cylinder}}$, $\mathrm{f_{zeppelin}}$, $\mathrm{S_{cylinder}}$ and $\mathrm{S_{zeppelin}}$ are the signal fractions and the diffusion weighted signal from the cardiomyocyte and extra-cellular components ($\mathrm{f_{cylinder}} + \mathrm{f_{zeppelin}} = 1$). While $\mathrm{R_{cylinder}}$, $\mathrm{D^{||}_{cylinder}}$, $\mathrm{D^{||}_{zeppelin}}$, and $\mathrm{D^{\perp}_{zeppelin}}$ are the cylinder radius and diffusivity and zeppelin diffusivities, respectively (Afzali et al., 2021).

The following parameters from the literature were used for the simulation (Farzi et al., 2021; Poole-Wilson, 1995; Tracy and Sander, 2011), $[\mathrm{f_{cylinder}}, \mathrm{f_{zeppelin}}, \mathrm{R_{cylinder}}, \mathrm{D^{||}_{cylinder}}, \mathrm{D^{||}_{zeppelin}}, \mathrm{D^{\perp}_{zeppelin}}] = [0.65, 0.35, 9\,\mu\mathrm{m}, 3\,\mu\mathrm{m}^2/\mathrm{ms}, 1.5\,\mu\mathrm{m}^2/\mathrm{ms}, 1\,\mu\mathrm{m}^2/\mathrm{ms}]$. The timing of the diffusion encoding waveform is 17, 8, and 17 ms for the pre-, pause- and post-duration, respectively (Figure S1 ). Fitting the kurtosis tensor leads to mean kurtosis (MK) = 0.31, axial kurtosis (AK) = 0.26 and radial kurtosis (RK) = 0.33 which is close to the values we obtained in the in vivo analysis. By increasing the effective diffusion time, mean kurtosis and radial kurtosis increase slightly (Table S1 ).

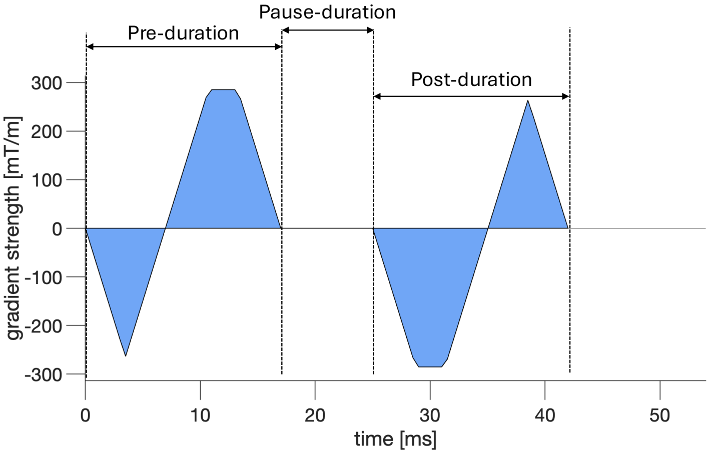


Figure S1 : Diffusion gradient waveform used in this study.

Table S1 : Calculated mean kurtosis (MK), axial and radial kurtosis (AK and RK) from simulated signal for different effective diffusion times.

| pre-pause-post duration [ms] | MK | AK | RK |
|---|---|---|---|
| 17-8-17 | 0.31 | 0.26 | 0.33 |
| 20-8-20 | 0.31 | 0.26 | 0.34 |
| 25-8-25 | 0.32 | 0.26 | 0.36 |
| 30-8-30 | 0.33 | 0.26 | 0.37 |
| 35-8-35 | 0.33 | 0.26 | 0.37 |

### References


Afzali, M., Nilsson, M., Palombo, M., Jones, D.K.. Spheriously? the challenges of estimating sphere radius non-invasively in the human brain from diffusion MRI. NeuroImage 2021;237:118183.

Farzi, M., Mcclymont, D., Whittington, H., Zdora, M.C., Khazin, L., Lygate, C.A., Rau, C., Dall'Armellina, E., Teh, I., Schneider, J.E.. Assessing myocardial microstructure with biophysical models of diffusion mri. IEEE transactions on medical imaging 2021;40(12):3775–3786.

Poole-Wilson, P.A.. The dimensions of human cardiac myocytes; confusion caused by methodology and pathology. Journal of molecular and cellular cardiology 1995;27(3):863–865.

Tracy, R.E., Sander, G.E.. Histologically measured cardiomyocyte hypertrophy correlates with body height as strongly as with body mass index. Cardiology research and practice 2011;2011(1):658958.
